# Dorsal Raphe Nucleus Enkephalin Peptide Modulates Behavioral Preference

**DOI:** 10.1101/2025.10.22.683947

**Authors:** Kathryn Braden, Andrew Trinagel, Eric Acevedo, Alexandra E. Bernstein, Marcela Arguello, Laura N. Massó-Quiñones, Aidan Evans-Strong, Samantha S. Dunn, Daniel C. Castro

## Abstract

The endogenous opioid system is a powerful modulator of motivation and affect. The dorsal raphe nucleus (DRN) in the midbrain has been established as an important site of opioid action and is an integral hub in behavioral modulation. To investigate the functional significance of DRN opioid signaling in aversive and appetitive behaviors we disrupted preproenkephalin (Penk) in DRN using CRISPR-Cas9 technology in Penk-Cre mice. We found that CRISPR mediated knockdown of enkephalin peptide in the DRN (DRN^Penk^) enhanced inflammation-induced mechanical sensitivity and odor avoidance. Additionally, loss of DRN^Penk^ diminished sucrose preference and engagement with a novel social stimulus. To further characterize the opioid system within the DRN, we performed Hiplex in situ hybridization of 12 genes in the same tissue. This revealed that DRN^Penk^ is largely separate from DRN serotonin cells and is instead distributed on glutamatergic and GABAergic cells. However, subtype-specific knockdown of DRN^Penk^ from glutamatergic and GABAergic cells did not replicate the behavioral effects of general DRN^Penk^ knockdown. This suggests that these neurons represent a novel population that mediate motivated behaviors distinctly from canonical DRN mechanisms.

## Introduction

The opioid epidemic will be remembered as one of the most shocking public health failures of the early 21^st^ century, leading to the deaths of over 500,000 people in the United States between 1999-2019 [1]. One major difficulty in combatting the opioid crisis arises from the fact that while opioids have a high propensity for abuse, they are still the single most effective acute analgesic available. Although significant investment into both the addictive and analgesic properties of opioids has increased in recent years, other, potentially life-changing, medical domains of the endogenous opioid system have gone under-appreciated. One example of this is the role endogenous opioids play in affective neuropsychiatric disorders such as depression or post-traumatic stress disorder (PTSD) [2–4]. Reaching a deeper understanding of how endogenous opioids contribute to behavioral and psychiatric disorders will help uncover the complete pharmacological potential of this system that is integral to brain function.

The dorsal raphe nucleus (DRN) is directly ventral to the periaqueductal gray nucleus (PAG) in the dorsal midbrain and is most well-known for its extensive serotonergic projections to the forebrain and its role in mood regulation. Though clearly important, these 5-hydroxytryptophan (5-HT) containing neurons only account for about 30% of all DRN cells, leaving the role of other neurotransmitter systems in this region unresolved [5]. Many of these non-serotonergic cells are enriched in opioid peptides, including enkephalin [6,7]. Enkephalin is encoded by the preproenkephalin (Penk) gene and is subsequently cleaved into the functional peptides met-enkephalin and leu-enkephalin [8,9]. Enkephalin peptides can bind and activate both delta (DOR) and mu opioid receptors (MOR), both of which are associated with mood, affect, motivation, and anxiety. [10–12]. Considering the wide-ranging interconnectedness of the DRN and the importance of endogenous opioids to behavioral tuning, it is likely that enkephalin originating from DRN neurons have a variety of behavioral effects beyond analgesia and reward. Due to technological constraints, studies investigating opioids in DRN have relied on receptor-level pharmacology to infer how endogenous peptides may contribute to behavioral effects [13–23]. However, whether these or other behavioral effects are based on the endogenous ligand availability has not been directly studied.

Here, we investigated how enkephalin peptide from the DRN (DRN^Penk^) contributes to behaviors relating to aversion and reward. Through a Cre-dependent CRISPR-Cas9 approach, we specifically knocked down DRN^Penk^ transcription and assessed behavioral differences in Penk-Cre+ mice and their Penk-Cre-controls. Additionally, we mapped the spatial distribution of the opioid system along with other neurotransmitters throughout the DRN through Hiplex in situ hybridization. Using this information, we also evaluated whether the behavioral effects of DRN^Penk^ knockdown were driven by a subpopulation of cells within the DRN.

## Materials and Methods

### Animals

Adult (18-35g) male and female C57BL/6J, preproenkephalin-IRES-Cre (*Penk*-Cre), Slc17a8-IRES2-Cre (*Vglut3*-Cre), Slc17a6-IRES-Cre (*Vglut2*-Cre), and Slc32a1-IRES-Cre (*Vgat*-Cre) were group housed and maintained on a 12-hour light-dark cycle. Mice were either bred at Washington University in Saint Louis or purchased from Jackson Laboratories (Bar Harbor, ME). Unless otherwise noted, mice had ad libitum access to food and water. All mice were kept in a sound-attenuated, isolated holding facility at least one week prior to surgery, post-surgery, and throughout the duration of behavioral assays to minimize stress. For Penk-Cre mice, we used Cre-cage and littermate controls. For Vglut2, Vglut3, and Vgat-Cre mice the cohorts were run in conjunction with Penk-Cre cohorts containing Penk-Cre-controls and following placement verification a group of vGlut2-Cre, vGlut3-Cre, or vGat-Cre (vG-Cre) mice that did not display viral expression in the DRN were created post-hoc as a “miss” group for comparison. There were no statistical differences between vG-Cre miss and Penk-Cre-behaviors so they were combined into one control group. Order of behavioral tests was counterbalanced across cohorts. All animals were monitored for health status daily before experimentation and for the entirety of the study. All procedures were approved by the Animal Care and Use Committee of Washington University in St. Louis and conformed to the NIH guidelines.

### Stereotaxic Surgery

Mice were anaesthetized in an induction chamber (2-5% isoflurane) and placed into a stereotaxic frame (Stoelting, Chicago, IL) and maintained on 1-2% isoflurane throughout the surgery. Viral injection in the DRN (AP -4.6, ML 0, DV -3.0) [24] of 100nL of a 2:1 dilution of AAV5-CMV-FLEX-SaCas9-U6-sgPenk (The Hope Center Viral Core-Washington University in St. Louis, MO): AAV1-Ef1a-DIO-EYFP (Addgene, Watertown, MA) was delivered with a Neuros needle syringe (Hamilton Company, Franklin, MA) at a rate of 50nL/min. Mice were allowed to recover for 4 weeks before any behavioral experiments to ensure robust viral expression.

### Behaviors

#### Carrageenan-induced mechanical sensitivity

Mice were placed on an elevated wire mesh rack underneath plexiglass boxes and allowed to habituate for one hour for 2 days prior to measuring baseline mechanical sensitivity by applying calibrated von Frey filaments (Stoetling; ranging from 3.22 (0.16g) to 4.31 (2.0g) bending force) to the plantar surface of the hind paw according to the Dixon up-and-down method [25]. Paw withdrawal threshold (PWT) was defined as the median 50% threshold force in grams needed to elicit a withdrawal response, calculated according to the Chaplan method [26]. Following baseline testing, mice were briefly placed under isoflurane (2%) anesthesia and 20μL of 0.1% λ-carrageenan (Sigma-Aldrich, St. Louis, MO) was injected subcutaneously into the plantar surface of the left hind paw. Mice were allowed to habituate to the mesh rack for at least one hour before mechanical sensitivity was re-assessed 3 hours following carrageenan injection.

#### Odor Preference

Odor preference was performed based on Witt et al. [27]. Before testing mice were placed in an empty habituation cage inside a biosafety cabinet for 15 minutes before being placed in the empty test cage inside a second biosafety cabinet for another 15 minutes. Filter paper with 50μL of either distilled water or an odorant (attractive: Putrescine [PUT] or aversive: 2-isobutyl-thiazole [IBT] diluted in distilled water to 85mM [28,29]) was introduced to the cage and placed on the opposite side of the mouse’s position. Behavior was video recorded for 3 minutes for post-hoc scoring by a researcher blinded to the genotype and odorant. Each mouse was tested twice, once with water and once with an odorant (either attractive or aversive) and time spent sniffing the filter paper was normalized to water sniffing within subjects. Test days were 48 hours apart and order of odor presentation (water/odorant first/second) was counterbalanced. Behavioral scoring was performed using Observer XT 17 (Noldus, Leesburg, VA).

#### Elevated Zero Maze

The EZM was made of grey plastic, 144 cm in circumference, comprised of four 36 cm sections (two opened and two closed) and elevated 40 cm above the floor with a 5.5 cm wide path with a 0.5 cm lip on each open section. Room lighting was maintained at 10lux to promote exploratory behavior. Mice were positioned head-first into a closed arm and allowed to freely roam for 7 mins. Time spent in open or closed arms was recorded and analyzed using Ethovision XT 17 (Noldus).

#### Social Interaction

Social Interaction was assayed based on Yang, et al. [30]. Briefly, mice were habituated to a three-chamber apparatus with free access to all three chambers for 10 minutes. Test mice were then removed from the apparatus and an age and sex-matched stranger mouse was placed underneath one pencil cup in the corner of one chamber and an identical pencil cup was placed empty upside down in the corner of the opposite chamber (placement randomized). Test mice were then placed into the center chamber and allowed to roam freely and behavior was recorded for 10 minutes. Time spent in each chamber throughout the tests was recorded and analyzed using Ethovision XT 17.

#### Sucrose Preference

Sucrose preference was assessed using a home cage two-bottle choice assay. Mice were singly housed and had ad libitum access to chow and two 15mL bottles. On days 1-3 one bottle was filled with water and one was left empty. Each day bottles were weighed and switched sides to promote sampling of both bottles, preventing development of a side bias. On days 4-5 the second bottle was filled with 1% sucrose solution and bottles were weighed and switched places after 24 hours. Sucrose preference was calculated as the amount of sucrose intake/total liquid intake × 100 during each test day and final sucrose preference scores are expressed as an average of the two test days.

#### Food Intake

Mice were habituated to plexiglass 25cm x 25cm chambers containing a Feeding Experimentation Device (FED) [31,32] filled with sucrose pellets (20mg) on a free-feeding setting. Habituation was performed for one hour for 3 days before mice were either food deprived or left ad libitum (order counterbalanced) for 24 hours. Then mice were placed back into the chamber and the number of sucrose pellets was recorded by the FED. Mice were then given the opposite treatment (food deprivation or ad libitum for 24 hours), and the test was repeated the final day.

### Tissue Processing

Animals were transcardially perfused with 0.1 M phosphate-buffered saline (PBS) and then 4% paraformaldehyde (PFA). Brains were dissected and post-fixed in 4% PFA overnight and transferred to 30% sucrose solution for cryoprotection. For histological analysis, brains were sectioned at 30 μM on a cryostat and thaw-mounted onto SuperFrost Plus slides (ThermoFisher, St.Louis, MO), then counterstained with Vectashield with DAPI (Vector Laboratories, Newark, CA), and coverslipped.

### RNAscope Fluorescent In Situ Hybridization

Fluorescent in situ hybridization was performed according to the RNAscope 2.0 Fluorescent Multiplex or Hiplex Kits’ User Manual for Fixed Frozen Tissue (Advanced Cell Diagnostics [ACD], Inc., Newark, CA). Following tissue processing as described above, brains were sectioned at 20 μM on a cryostat and thaw-mounted onto SuperFrost Plus slides. Slides were stored at −80°C until further processing.

Slides were post-fixed in 4% PFA, dehydrated, and treated with pretreatment protease III solution according to the manufacturer’s protocol. Hiplex slides were then incubated with target probes for mouse proenkephalin (*Penk*, NM_001002927.2, probe region 106 - 1332), prepronociceptin (*Pnoc*, NM_010932.2, probe region 325 - 1263), prodynorphin (*Pdyn*, NM_018863.3, probe region 33-700), mu opioid receptor (*Oprm1*, NM_001039652.1, probe region 1135 - 2162), delta opioid receptor (*Oprd1*, NM_013622.3, probe region 1214 - 2492), kappa receptor (*Oprk1*, NM_001204371.1, probe region 256-1457), nociception receptor (*Oprl1*, NM_011012.5, probe region 988 - 1937), vesicular GABA transporter (*slc32a1*, NM_009508.2, probe region 894 – 2037), vesicular glutamate transporter type 2 (*slc17a6*, NM_080853.3, probe region 1986 - 2998), vesicular glutamate transporter type 3 (*slc17a8*, NM_182959.3, probe region 781 - 1695), tryptophan hydroxylase 2 (*Tph2*, NM_173391.3, probe region 1640 - 2622), and tachykinin precursor 1 (*Tac1*, NM_009311.2, probe region 20 - 1034). All target probes except Pdyn and Tac1 consisted of 20 ZZ oligonucleotides and all probes were obtained from ACD. Following probe hybridization, sections underwent a series of probe signal amplification steps followed by incubation of three fluorescently labeled probes designed to target the specific channel associated with the probes. Slides were counterstained with DAPI, and coverslips were mounted with Prolong Gold Antifade Mountant (ThermoFisher). All images were obtained on an Echo Revolution Hybrid Microscope (Discover Echo Inc., San Diego, CA) and analyzed with HALO software. Following imaging of the first three sets of probes on the Hiplex slides, fluorophore cleavage was performed and the next set of fluorophores were hybridized. This process was repeated four times to label all 12 probes in the same slices. The DAPI layers for each 4 hybridization sets were registered using the Hiplex Image Registration Software (ACD) and registered each channel image resulting in a single 12-channel image. These images were opened in Halo Software (Indica Labs, Albuquerque, NM) and boundaries were set for the area to be analyzed. DAPI positive cells were then registered and used as markers for individual cells. The maximum area around each cell for probes to be detected was then set, approximately 6 microns.

### Statistical Analysis

All data collected were averaged and expressed as mean ± SEM. Statistical significance was taken as *p < 0.05, **p < 0.01, ***p < 0.001, and ****<0.0001 as determined by Student’s t test, one-way, or two-way repeated-measures ANOVA as appropriate. For behavioral experiments we used one-way or two-way repeated-measures ANOVA followed by a Tukey or Sidak post hoc tests. All n-values for each experimental group are described in the appropriate figure legend. Statistical tests did not detect any significant differences between male and female mice and were therefore combined to complete final group sizes. For behavioral experiments, group size ranged from n = 8-20. For *in situ* hybridization quantification experiments, slices were collected from 3 female mice, with data averaged from 3 slices per mouse. Statistical analyses were performed in GraphPad Prism 8.0 (Graphpad, La Jolla, CA).

## Results

### Knockdown of DRN enkephalin enhances aversive behaviors

We injected a previously characterized viral vector containing Cre-dependent CRISPR-Cas9 directed to the Penk locus as well as an eYFP fluorescent marker [6] into the DRN of Penk-Cre+ mice and their littermate Cre-controls (Fig 1A). This led to expression of eYFP restricted to the DRN (Fig 1B and 1C left) and resulted in a 50% reduction in Penk gene expression in Penk-Cre+ mice compared to controls, as measured by fluorescent in situ hybridization (Fig 1C right and 1D), but had no effect on Pdyn gene expression (Fig 1C right and 1E). Following 4 weeks of recovery and incubation of the virus, mice underwent behavioral testing to assess how knockdown of DRN^Penk^ affected aversive and appetitive responses.

**Figure 1:**
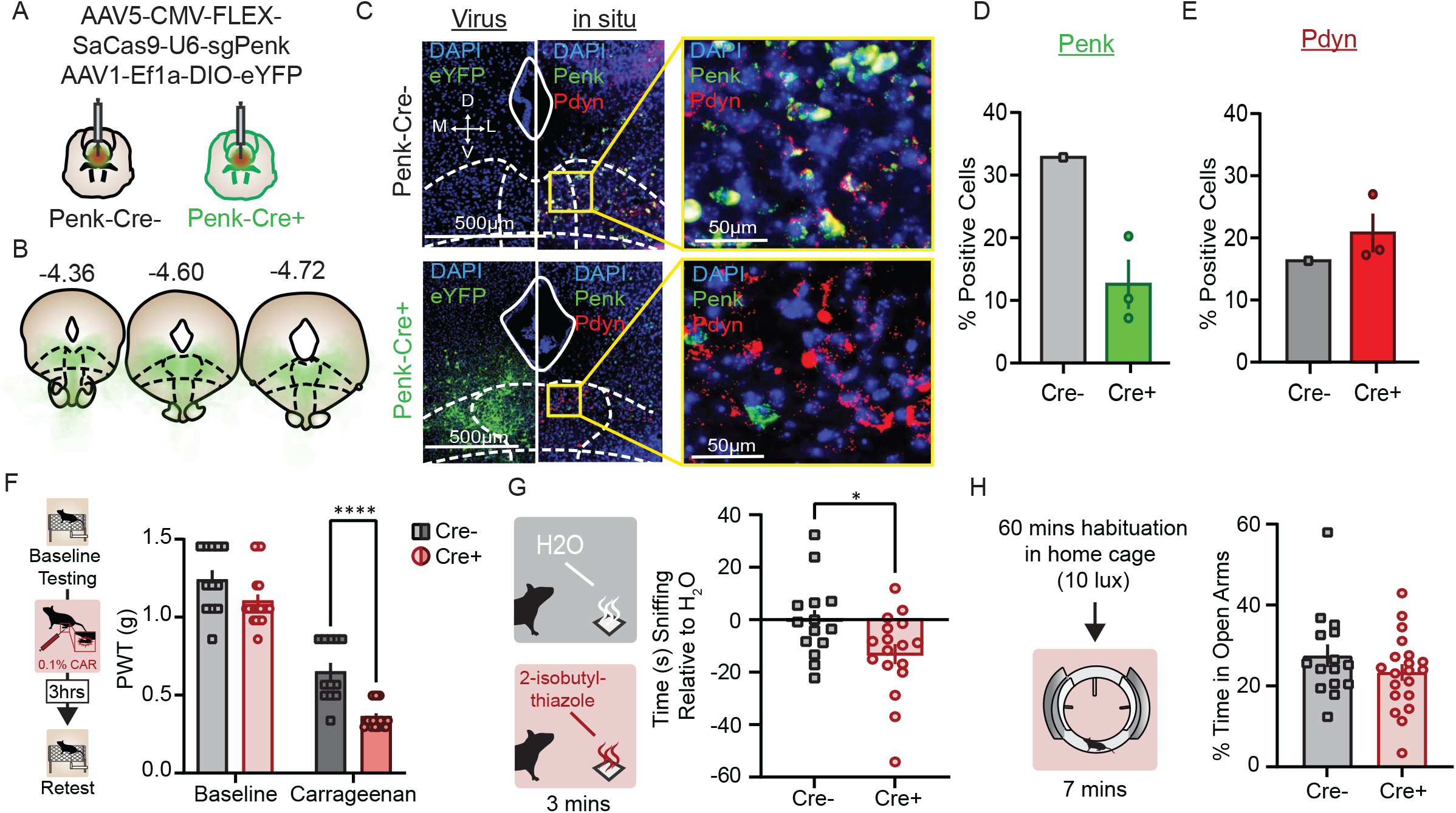
DRN^Penk^ knockdown enhances aversion. A) Schematic of knockdown strategy B) Map of viral spread throughout DRN of mice included in behavioral analysis C) Representative image of viral spread (left) and in situ hybridization (right) for *Penk* and *Pdyn* in Penk-Cre-(top) and Penk-Cre+ (bottom) DRN. Yellow boxes outline zoomed in region in panels on right. Images were brightened for display purposes, but analysis was run on original unedited images. D-E) Quantification of *Penk* (D) and *Pdyn* (E) FISH signal as % of cells within DRN positive for each transcript, Cre+ n= 3 mice, 2-3 slices per mouse; Cre-n=1 mouse, 2 slices. E) Mechanical sensitivity as paw withdrawal threshold (PWT) in grams of mice before (Baseline) or 3 hours after (Carrageenan) intraplantar carrageenan (0.1% in saline) injection, Cre+ n=19; Cre-n= 12. Cre+ mice had enhanced mechanical allodynia post carrageenan injection, Two-Way ANOVA with Sidak’s multiple comparison;****p<0.0001 F) Time sniffing IBT infused filter paper relative to time sniffing water infused filter paper during 3 minute test, Cre-n=14; Cre+ n=17. Cre+ mice avoided IBT filter paper more than Cre-mice, Two-Way ANOVA with Sidak’s multiple comparison; *p=0.0268 G) Percent of time spent in open arms during EZM test for Cre- and Cre+ mice, Cre+ n= 20; Cre-n= 15. Data are all mean ± SEM.

First, we assessed how loss of DRN^Penk^ affected mechanical sensitivity, as dorsal midbrain enkephalin is canonically associated with descending pain modulation [33]. Knockdown of enkephalin did not lead to significant changes in paw withdrawal thresholds (PWT) at baseline (Fig. 1F left). However, following an intraplantar injection of low-dose carrageenan to induce a mild inflammatory reaction, Penk-Cre+ mice had a hyperalgesic response compared to Penk-Cre-controls (Fig. 1F right). Apart from pain, opioid activity in the midbrain has also been implicated in aversive responses [34,35]. Mice display robust behavioral responses to odors, and their natural behaviors are highly influenced by olfactory cues. With this in mind, we chose to use odor avoidance to gauge aversion independent of nociception or fear. We used the aversive odorant, 2-isobutyl-thiazole (IBT) [28,29] in an odor preference assay and normalized individual sniffing behavior to a neutral odor (water) within subjects. Compared to Cre-controls, Penk-Cre+ mice avoided IBT-infused filter paper more than water (Fig. 1G, Supplementary Fig. 1), suggesting an enhanced aversion. Although IBT has previously been shown to be aversive to mice, our Cre-control mice were on average interacting with IBT filter paper the same amount as water filter paper. This was likely driven by an order effect in our counterbalanced presentation of either H2O or IBT (Supplementary Fig. 1G-H). The presentation of IBT first led to a decreased sniffing time of water during the second presentation in Cre-controls (Fig. 1G) which resulted in a neutral relative preference for IBT over water (Supplementary Fig. 1H), however in either condition (IBT presented first or second) Penk-Cre+ mice sniffed IBT filter paper less than controls (Supplementary Fig. 1G-H). However, there were no differences between Penk-Cre+ and Cre-controls in general anxiety levels, as measured by an EZM assay (Fig. 1H).

### Knockdown of DRN enkephalin impairs appetitive behaviors

Opioids throughout the brain also impact rewarding and appetitive behaviors, so we assessed a battery of appetitive assays with our Penk-Cre+ and Cre-mice to determine how DRN^Penk^ influences the opposite valence. In the first 3 minutes of a social interaction test, Penk-Cre-controls spent more time investigating a novel mouse than a novel object whereas Penk-Cre+ mice did not display this preference (Fig. 2A, Supplementary Fig. 2A-D), suggesting the appetitive nature of the social stimulus may have been diminished with knockdown of DRN^Penk^. This effect was not due to differences in locomotion, as total distance traveled was similar between genotypes (Supplementary Fig. 2E). Similarly, preference for a palatable sucrose solution (1%) over water was decreased in Penk-Cre+ mice (Fig. 2B) by 15% compared to controls, but total liquid consumed did not differ (Supplementary Fig. 2F). In an odor preference assay using the normally appetitive odorant, putrescine (PUT) [28], Cre-mice spent more time sniffing the PUT-infused filter paper than water, however Penk-Cre+ mice had a neutral preference, spending on average the same amount of time investigating either PUT and water filter papers (Fig. 2C, Supplementary Fig. 3A-D). Similarly to the IBT test, this effect may have been diluted by our counterbalanced approach: The effect of PUT preference over water in the Cre-mice was stronger when PUT was presented before the water test (Supplementary Fig. 3E-F). Regardless, in both tests the Penk-Cre+ mice appear to have a relatively lower preference compared to Cre-mice. Finally, the preference impairment observed in Cre+ mice did not extend to general food intake or reward as sucrose pellet intake during either ad libitum or food deprived states were similar in both genotypes (Fig. 2D).

**Figure 2:**
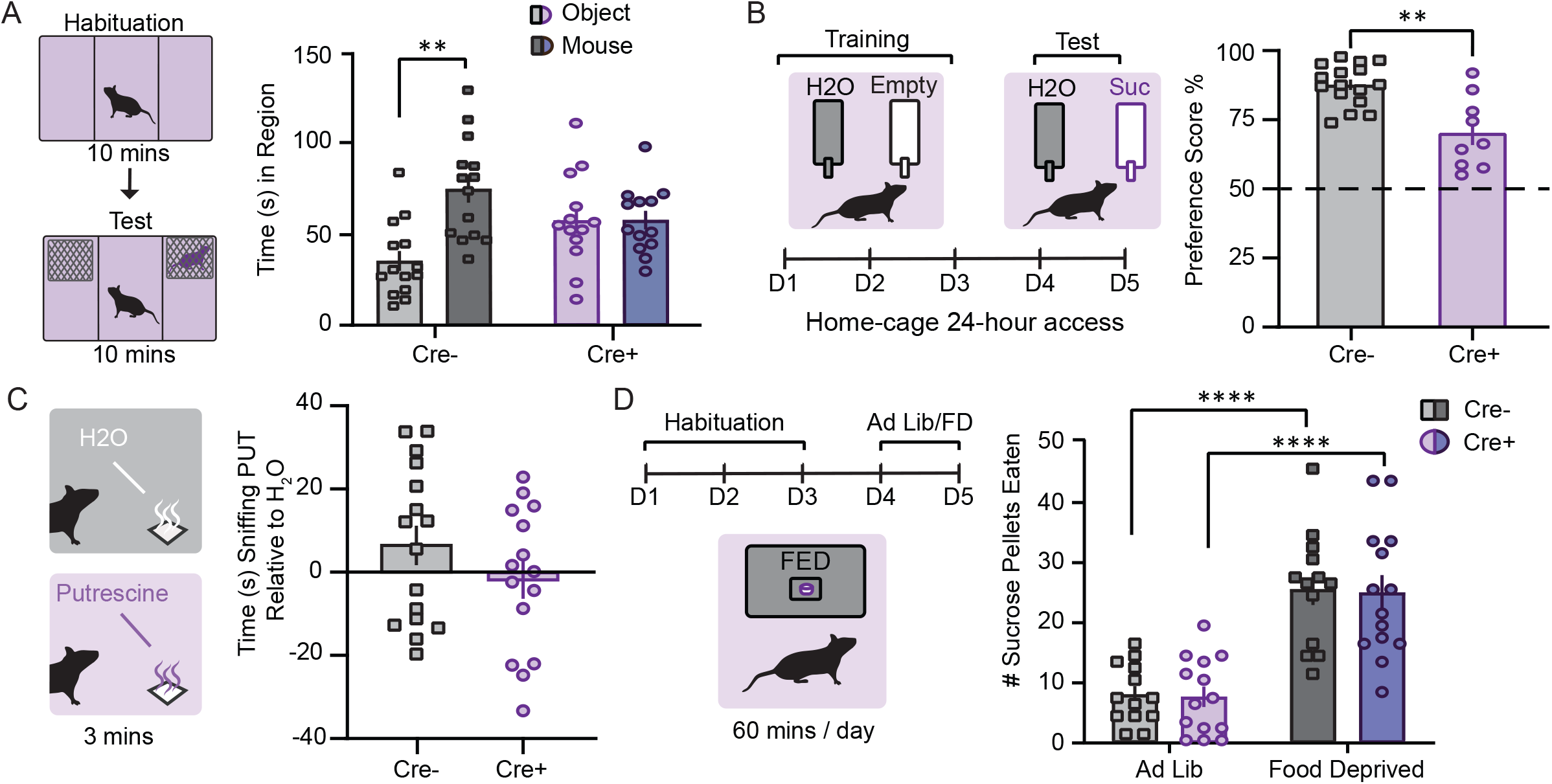
DRN^Penk^ knockdown impairs behavioral preference. A) Social interaction of Penk-Cre+ or Cre-controls. Cre-mice spend more time in the region containing a novel mouse than a novel object, while Cre+ mice spend similar times in both regions after the initial 3 minutes of testing. Cre-n = 14; Cre+ n=13, Two-Way ANOVA with Sidak’s multiple comparison Mouse vs. Object; **p= 0.0025 B) Sucrose Preference. Preference Score calculated by the equation (Sucrose consumed/Total consumed)*100 for Cre- and Cre+ mice, Cre+ mice have a lower preference for sucrose than Cre-controls. Cre-n= 16 Cre+ n=9, Unpaired t-test with Welch’s correction; **p=0.0037. C) In an odor preference assay Cre-mice spend more time sniffing PUT infused filter paper relative to time sniffing water infused filter paper during 3 minute test, Cre-n=12; Cre+ n=15. D) No differences in sucrose intake between Penk-Cre+ or Cre-mice; Total # sucrose pellets eaten during either ad libitum (Ad lib) or food deprived (FD) states for Cre- and Cre+ mice, Cre+ n= 14; Cre-n= 13, Two-Way ANOVA with Sidak’s multiple comparison Ad Lib vs. FD; ****p<0.0001. Data are all mean ± SEM.

### Landscape of endogenous opioid system within the DRN

The DRN has been molecularly characterized multiple times, but most of these efforts were strictly defined by the expression of serotonin [36–38]. As discussed in multiple publications [5,6,39], the serotonin cells only account for ∼20-30% of all neurons within the DRN, leaving unresolved how other cell types may be colocalized or spatially distributed. To get a complete picture of how the endogenous opioid system within this region is composed, we performed Hiplex fluorescent in situ hybridization (FISH) of 12 genes, including the opioid peptides and receptors (Fig. 3 and Table 1). We found that all components of the opioid system are expressed in the DRN and overlap with other major neurotransmitter systems (Fig. 3A-I and Table 1). The *Penk* gene specifically is expressed in approximately 25% of all cells in this region (Fig 3B, H, I and Table 1). Although the DRN is usually synonymous with 5-HT, we found that only about 20% of cells in this region expressed the serotonergic cell marker *Tph2* (Table 1; Fig. 3D, H-K) and these were largely non-overlapping with the *Penk*+ cells (Table 1; Fig 3K). Instead *Penk*+ cells in this region were co-expressed with either glutamatergic or GABAergic cell markers (Table 1; Fig. 3J). The DRN cells that co-express *Penk* and *Tph2* are mostly triple labeled with vGlut3 (Fig. 3K-L). This suggests that DRN^Penk^ is a unique population of cells that have not yet been characterized.

**Figure 3:**
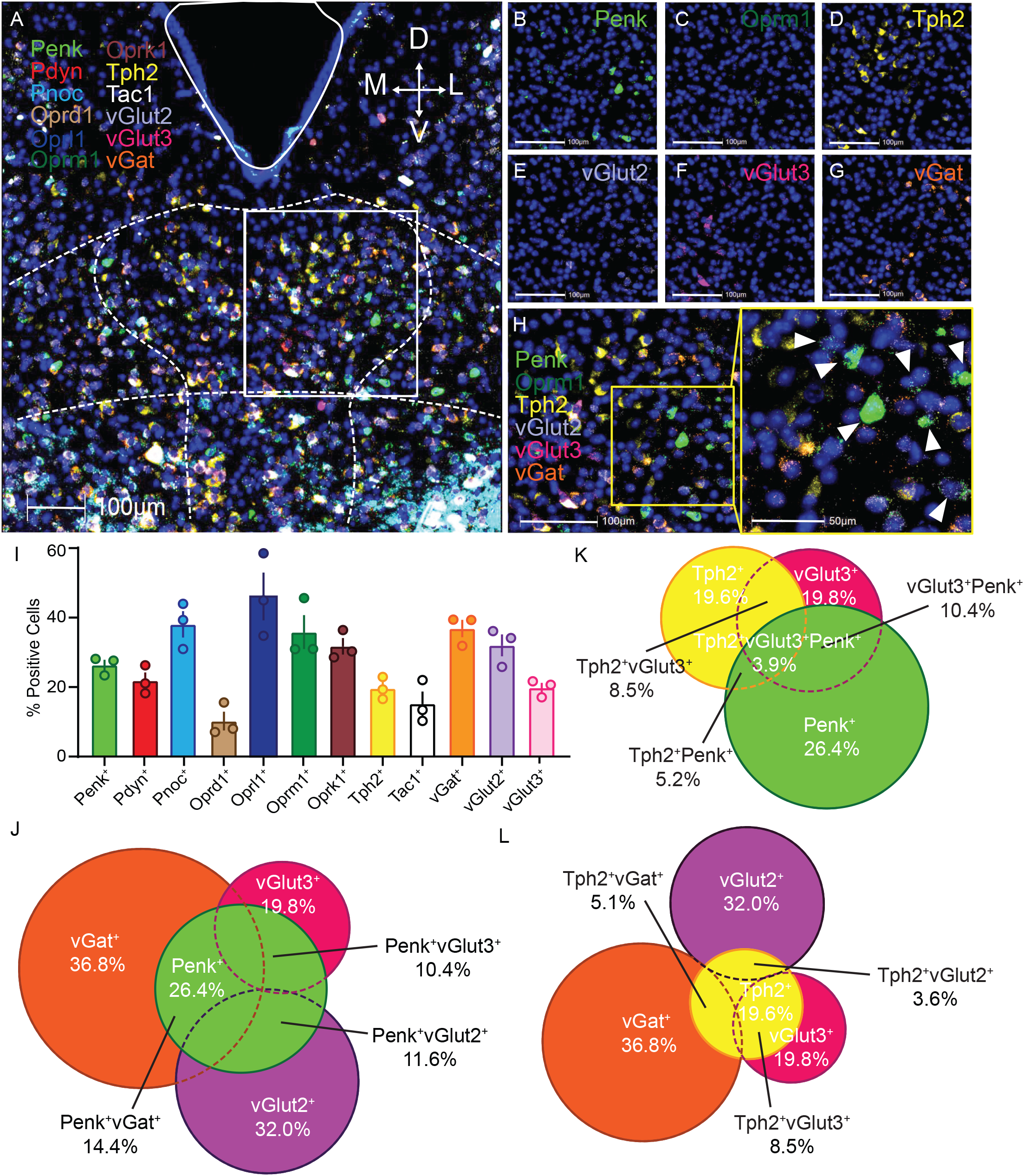
Hiplex Fluorescent in situ hybridization (FISH) of DRN. A) Representative image of In situ hybridization of *Penk, Pdyn, Pnoc, Oprd1, Oprl1, Oprm1, Oprk1, Tph2, Tac1, Slc17a6, Slc17a8, Slc32a1* in DRN (scale bar= 100μm). B-G) Zoomed in and channel separated images of the white square in A of *Penk* (B), *Oprm1* (C), *Tph2* (D), *Slc17a6* (E), *Slc17a8* (F), *Slc32a1* (G). H) Merged image of B-G channel images. Yellow square shows zoomed in image (right) with Penk labeled cells indicated by white arrows. Images were brightened for display purposes, but analysis was run on original unedited images. I) Quantification of FISH signal as % of cells within DRN + for each transcript, Data are mean ± SEM, n=3 mice, 3 slices per mouse. J) Quantification of FISH depicting overlap of Penk+ and Tph2+ cells within the DRN. K) Quantification of FISH depicting overlap of Tph2+, Slc32a1+, Slc17a6+, and Slc17a8+ cells within the DRN. L) Quantification of FISH depicting overlap of Penk+, Slc32a1+, Slc17a6+, and Slc17a8+ cells within the DRN.

**Table 1:**
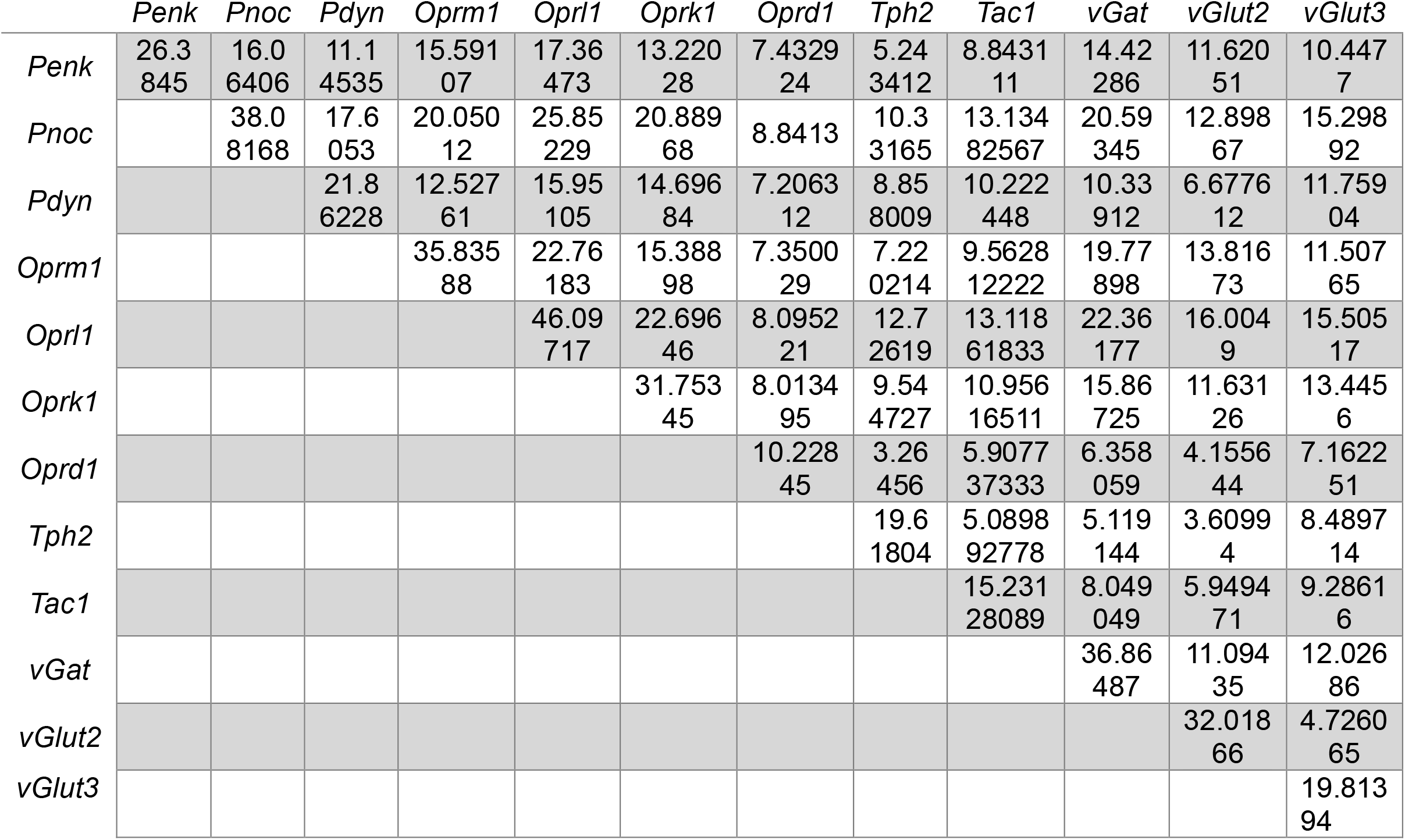
Expression and Overlap of Opioid System in DRN. Quantification of FISH signal as % of cells within DRN + for each probe and the corresponding overlap with each other probe used. Data are mean, n= 3 female mice, n= 3 slices per mouse.

### Effects of enkephalin knockdown are not driven by glutamatergic or GABAergic subpopulations

Neuropeptides, such as opioids, are commonly co-released by neurons along with other neurotransmitters such as GABA and glutamate to fine-tune synaptic communication [40,41]. Previous literature indicates DRN MOR activation interacts with GABAergic interneurons to disinhibit serotonin efflux [42–44]. This interaction has been postulated to contribute to other behavioral effects of DRN opioid signaling such as fear [45], anxiety [44], reward [39], and depression [15]. To determine whether the behavioral effects we observed from DRN^Penk^ knockdown were driven by co-release with either GABA or glutamate, we used vGat-, vGlut2-, and vGlut3-Cre mice to selectively knockdown enkephalin in each glutamatergic or GABAergic subpopulation of the DRN (Fig. 4A) and ran the same behaviors to see if we could replicate our original phenotype of DRN^Penk^ knockdown. A control group consisting of concurrently run Penk-Cre-mice and vG-Cre+ mice with no viral expression were used for comparison (Fig 4A, top left). In social interaction, control mice preferred the novel mouse over the novel object (Fig. 4B, Supplementary Fig. 4A, E), as expected, but interestingly the knockdown of DRN^Penk^ from any of the vG-Cre subtypes was sufficient to dampen this preference (Fig. 4B, Supplementary Fig. 4B-D, F-H) but did not affect locomotion (Supplementary Fig. 4I). The preference for mouse over object was most preserved in the vGlut2 knockdowns, but both vGat and vGlut3 knockdown nearly abolished any preference (Supplementary Fig. 4F-H). However, when it comes to odor avoidance, all iterations of DRN^Penk^ knockdown avoided IBT-infused filter paper more than water-infused paper, but there was no significant difference when compared to the control mice run at the same time (Fig 4B). This cohort of control mice demonstrated a more pronounced aversion to IBT than the cohorts in Fig. 1G but their behavior was in the same ranges of sniffing time and interactions and had a similar ordering effect (Supplementary Fig. 5A-E). Importantly, the vG-Cre cohorts run at the same time were not significantly different from the controls, suggesting the aversion was not enhanced by the knockdown of DRN^Penk^ from these neuronal subtypes as observed in Penk-Cre+ mice (Supplementary Fig. 5). Additionally, the knockdown of DRN^Penk^ in only vGlut2, vGlut3, or vGat cells was not sufficient to replicate the carrageenan-induced hypersensitivity (Fig. 4D). There was also no effect on sucrose preference (Fig. 4E), in contrast to general targeting of DRN^Penk^ (Fig 1-2). In sum, no subtype of DRN^Penk^ knockdown fully recapitulated the enhanced aversion and diminished preference of general DRN^Penk^ knockdown, but all of them had some effects on social interaction but with varying degrees of success. This suggests these effects as a whole are specific to the opioid peptide, DRN^Penk^, rather than classic neurotransmitters such as glutamate, serotonin, or GABA.

**Figure 4:**
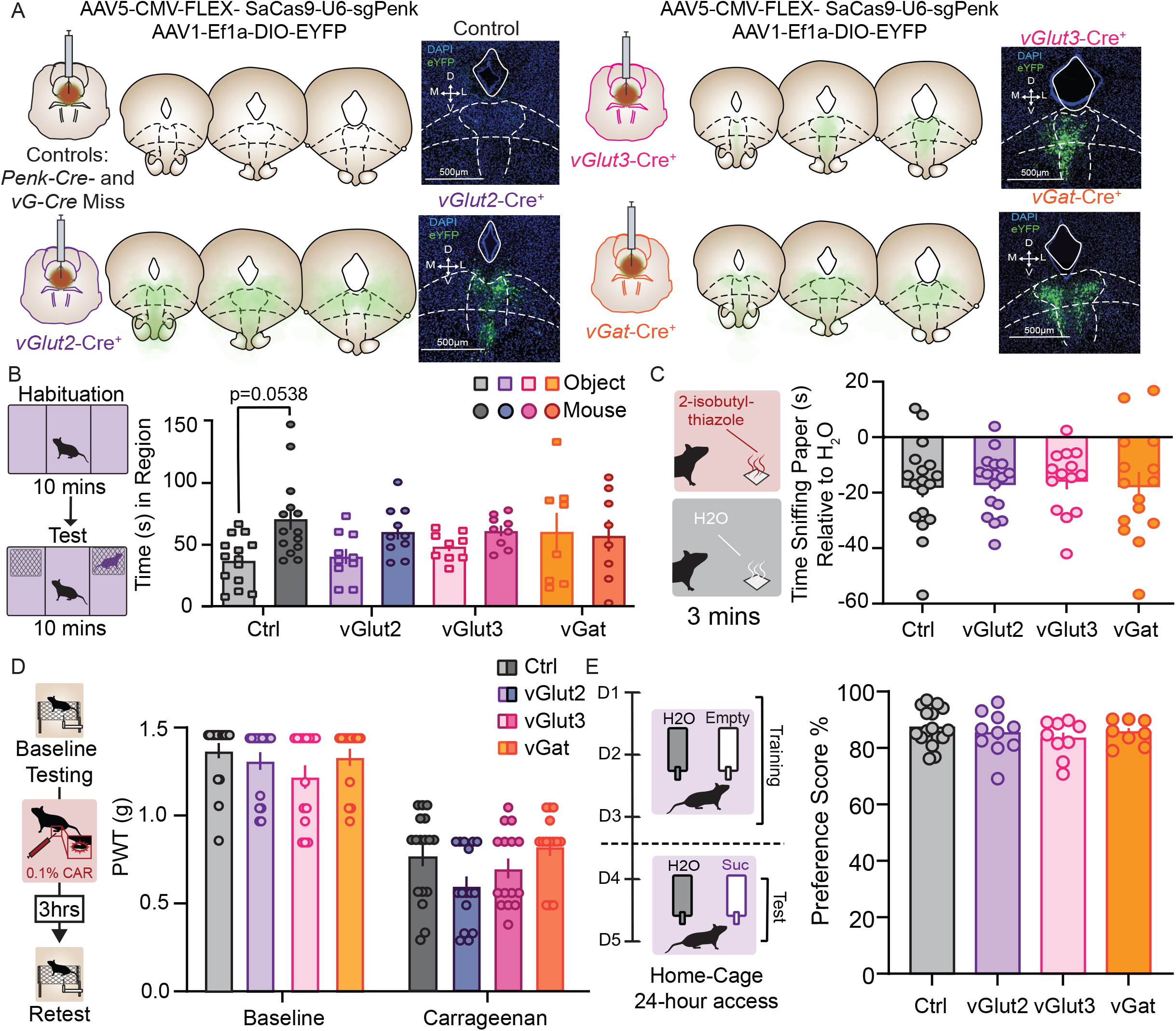
Behavioral effects of DRN^Penk^ knockdown are not driven by glutamatergic or GABAergic subpopulations. A) Schematic of knockdown strategy with maps of viral spread throughout DRN of mice included in behavioral analysis for each genotype and representative images of viral spread. Images were brightened for display purposes. B) Time spent in either the Object or Mouse region during the first 3 minutes of a social interaction test of Control and vG-Cre mice. Control mice trend toward spending more time in the region containing a mouse than a novel object, Two-Way ANOVA with Sidak’s multiple comparison Mouse vs. Object; Control p=0.0538. Control n=14 (n=8 Cre-, n=6 vG-Cre miss), vGlut2 n=11, vGlut3 n=9, vGat n=8 C) Time sniffing IBT infused filter paper relative to time sniffing water infused filter paper during 3 minute test, Control n= 18 (n=10 Cre-, n=8 vG-Cre miss), vGlut2 n= 17, vGlut3 n= 14, vGat n= 14. All genotypes avoided the IBT filter paper more than water and there were no differences between genotypes. D) Mechanical sensitivity as paw withdrawal threshold (PWT) in grams of mice before (Baseline) or 3 hours after (Carrageenan) intraplantar carrageenan (0.1% in saline) injection. All genotypes had a moderate reduction in PWT after carrageenan. Control n= 17 (n=9 Cre-, n=8 vG-Cre miss), vGlut2-Cre n= 17, vGlut3-Cre n= 14, vGat-Cre n= 14 E) Sucrose Preference Score calculated by the equation (Sucrose consumed/Total consumed)*100 was similar in all genotypes. Control n= 16 (n=10 Cre-, n=6 vG-Cre miss), vGlut2-Cre n= 10, vGlut3-Cre n= 9, vGat-Cre n= 8. Data are all mean ± SEM.

## Discussion

In these studies, we sought to understand how DRN^Penk^ neurons contribute to the processing of appetitive and aversive behavior. We found that deletion of enkephalin from DRN cells led to an overall enhancement of aversion (e.g. inflammatory pain and IBT odorant) and suppression of appetitive stimuli (e.g. social interaction and sucrose water). However, DRN^Penk^ did not appear to mediate overt motivation or non-specific anxiety, as time in the open arms in EZM and sucrose pellet consumption were unaffected by DRN^Penk^ deletion. This implies that DRN^Penk^ deletion does not generate aversion per se but instead shifts the preference for salient stimuli towards an aversive bias. This bias resembles that of neuropsychiatric disorders where aversion is enhanced, and reward is diminished such as chronic pain, PTSD, or addiction. Future studies will investigate how DRN^Penk^ is affected by disease states and whether this population of neurons may be targeted for therapeutic potential.

DRN is generally defined by its serotonergic neurons. However, immunohistochemistry, ISH, and single-cell RNA-seq experiments all indicate that DRN is comprised of a rich constellation of neuropeptides and transmitters that influence affective and motivated behaviors. Using Hiplex FISH, we mapped the spatial composition of the opioid family within DRN, revealing that the enkephalin system is largely independent from the serotonergic system. But similar to other published reports, we found that both glutamatergic and GABAergic cells within the DRN overlapped with the opioid peptides (including enkephalin) and receptors [6,39,46]. Contrary to our hypothesis, the behavioral effects of enkephalin knockdown did not depend on the specific GABAergic or glutamatergic subpopulations; rather, the behavioral effects were driven by the relative proportion of enkephalin deleted from the DRN and the sensitivity of the assay. For example, social interaction was modulated by knockdown in vGlut2-, vGlut3-, and vGat-Cre subpopulations. In contrast, inflammatory hypersensitivity, odor avoidance, and sucrose preference were unaffected. One potential explanation could be that social interaction is more sensorily complex relative to odor, mechanical sensitivity, and taste. Thus, less extensive dysregulation of enkephalin was able to hamper multiple points of affective-sensory integration, thereby resulting in net disruption of behavior more similar to that of general knockdown. Regardless, these data suggest that DRN^Penk^ is a unique population of cells that regulates behavior and that those effects are based on a complex interplay of factors. Though not tested here, it also would be worthwhile to determine whether the effects observed extend to conditioned behaviors (e.g., operant learning, conditioned place preference), or whether it is restricted to unconditioned behaviors.

Since Penk coexpression in glutamatergic or GABAergic cell types was insufficient to drive behavioral changes, we hypothesize that the role of DRN^Penk^ may be related to its downstream outputs. Recent work has shown that one such enkephalin circuit from DRN to nucleus accumbens medial shell projection can enhance consummatory behaviors via MOR stimulation on efferent terminals. However, the enkephalin peptide itself was not involved in the consummatory phenotype, consistent with our results here, in which loss of enkephalin had no direct impact on overall consumption. Looking forward, assessing this projection or other circuit mechanisms may provide novel insights into the role of DRN^Penk^ on appetitive and aversive behaviors.

## Author Contributions

KB designed and performed experiments, collected and analyzed data, procured funding, and wrote the manuscript. AT, EA, AEB, MA, LNMQ, AES, and SSD performed experiments and collected data. AT and AEB helped edit the manuscript. DCC led the design and analysis and provided oversight of experiments, procured funding, and helped write and edit the manuscript.

All authors had final approval of the version to be published and agree to be accountable for all aspects of the work.

## References

1. Biancuzzi H, Dal Mas F, Brescia V, Campostrini S, Cascella M, Cuomo A, et al. Opioid Misuse: A Review of the Main Issues, Challenges, and Strategies. Int J Environ Res Public Health. 2022;19:11754.

2. Limoges A, Yarur HE, Tejeda HA. Dynorphin/kappa opioid receptor system regulation on amygdaloid circuitry: Implications for neuropsychiatric disorders. Front Syst Neurosci. 2022;16.

3. Callaghan CK, Rouine J, O’Mara SM. Chapter 3 - Potential roles for opioid receptors in motivation and major depressive disorder. In: O’Mara S, editor. Progress in Brain Research, vol. 239, Elsevier; 2018. p. 89–119.

4. Johnson BN, McKernan LC, Bruehl S. A Theoretical Endogenous Opioid Neurobiological Framework for Cooccurring Pain, Trauma, and Non-suicidal Self-injury. Curr Pain Headache Rep. 2022;26:405–414.

5. Braden K, Castro DC. The role of dorsal raphe nucleus neuropeptides in reward and aversion. Front Behav Neurosci. 2025;19.

6. Castro DC, Oswell CS, Zhang ET, Pedersen CE, Piantadosi SC, Rossi MA, et al. An endogenous opioid circuit determines state-dependent reward consumption. Nature. 2021;598:646–651.

7. Pomrenze MB, Cardozo Pinto DF, Neumann PA, Llorach P, Tucciarone JM, Morishita W, et al. Modulation of 5-HT release by dynorphin mediates social deficits during opioid withdrawal. Neuron. 2022. 5 October 2022. 10.1016/j.neuron.2022.09.024.

8. Rysztak LG, Jutkiewicz EM. The role of enkephalinergic systems in substance use disorders. Front Syst Neurosci. 2022;16.

9. Akil H, Watson SJ, Young E, Lewis ME, Khachaturian H, Walker JM. Endogenous opioids: biology and function. Annu Rev Neurosci. 1984;7:223–255.

10. Ragnauth A, Schuller A, Morgan M, Chan J, Ogawa S, Pintar J, et al. Female preproenkephalin-knockout mice display altered emotional responses. Proc Natl Acad Sci U S A. 2001;98:1958–1963.

11. Bérubé P, Poulin J-F, Laforest S, Drolet G. Enkephalin Knockdown in the Basolateral Amygdala Reproduces Vulnerable Anxiety-Like Responses to Chronic Unpredictable Stress. Neuropsychopharmacology. 2014;39:1159–1168.

12. Hayward MD, Pintar JE, Low MJ. Selective Reward Deficit in Mice Lacking β-Endorphin and Enkephalin. J Neurosci. 2002;22:8251–8258.

13. Bruchas MR, Schindler AG, Shankar H, Messinger DI, Miyatake M, Land BB, et al. Selective p38α MAPK deletion in serotonergic neurons produces stress resilience in models of depression and addiction. Neuron. 2011;71:498–511.

14. Land BB, Bruchas MR, Schattauer S, Giardino WJ, Aita M, Messinger D, et al. Activation of the kappa opioid receptor in the dorsal raphe nucleus mediates the aversive effects of stress and reinstates drug seeking. Proc Natl Acad Sci U S A. 2009;106:19168–19173.

15. Lutz P-E, Ayranci G, Chu-Sin-Chung P, Matifas A, Koebel P, Filliol D, et al. Distinct Mu, Delta, and Kappa Opioid Receptor Mechanisms Underlie Low Sociability and Depressive-Like Behaviors During Heroin Abstinence. Neuropsychopharmacol. 2014;39:2694–2705.

16. Nectow AR, Schneeberger M, Zhang H, Field BC, Renier N, Azevedo E, et al. Identification of a Brainstem Circuit Controlling Feeding. Cell. 2017;170:429–442.e11.

17. Schindler AG, Messinger DI, Smith JS, Shankar H, Gustin RM, Schattauer SS, et al. Stress produces aversion and potentiates cocaine reward by releasing endogenous dynorphins in the ventral striatum to locally stimulate serotonin reuptake. J Neurosci. 2012;32:17582–17596.

18. Wager TD, Scott DJ, Zubieta J-K. Placebo effects on human mu-opioid activity during pain. Proc Natl Acad Sci U S A. 2007;104:11056–11061.

19. Yu W, Pati D, Pina MM, Schmidt KT, Boyt KM, Hunker AC, et al. zPeriaqueductal Grey/Dorsal Raphe Dopamine Neurons Contribute to Sex Differences in Pain-Related Behaviors. Neuron. 2021;109:1365–1380.e5.

20. Campion KN, Saville KA, Morgan MM. Relative contribution of the dorsal raphe nucleus and ventrolateral periaqueductal gray to morphine antinociception and tolerance in the rat. Eur J Neurosci. 2016;44:2667– 2672.

21. Yaksh TL, Yeung JC, Rudy TA. Systematic examination in the rat of brain sites sensitive to the direct application of morphine: Observation of differential effects within the periaqueductal gray. Brain Research. 1976;114:83–103.

22. Ferreira MD, Menescal-de-Oliveira L. Nociceptive vocalization response in guinea pigs modulated by opioidergic, GABAergic and serotonergic neurotransmission in the dorsal raphe nucleus. Brain Research Bulletin. 2014;106:21–29.

23. Fardin V, Oliveras J-L, Besson J-M. A reinvestigation of the analgesic effects induced by stimulation of the periaqueductal gray matter in the rat. I. The production of behavioral side effects together with analgesia. Brain Research. 1984;306:105–123.

24. Franklin KBJ, Paxinos G. The Mouse Brain in Sterotaxic Coordinates. Third. Academic Press; 2007.

25. Dixon WJ. Staircase bioassay: The up-and-down method. Neuroscience & Biobehavioral Reviews. 1991;15:47–50.

26. Chaplan SR, Bach FW, Pogrel JW, Chung JM, Yaksh TL. Quantitative assessment of tactile allodynia in the rat paw. Journal of Neuroscience Methods. 1994;53:55–63.

27. M. Witt R, M. Galligan M, R. Despinoy J, Segal R. Olfactory Behavioral Testing in the Adult Mouse. J Vis Exp. 2009:949.

28. Saraiva LR, Kondoh K, Ye X, Yoon K, Hernandez M, Buck LB. Combinatorial effects of odorants on mouse behavior. Proceedings of the National Academy of Sciences of the United States of America. 2016;113:E3300.

29. Manoel D, Makhlouf M, Arayata CJ, Sathappan A, Da’as S, Abdelrahman D, et al. Deconstructing the mouse olfactory percept through an ethological atlas. Curr Biol. 2021;31:2809–2818.e3.

30. Yang M, Silverman JL, Crawley JN. Automated Three-Chambered Social Approach Task for Mice. Current Protocols in Neuroscience. 2011;56:8.26.1-8.26.16.

31. Matikainen-Ankney BA, Earnest T, Ali M, Casey E, Wang JG, Sutton AK, et al. An open-source device for measuring food intake and operant behavior in rodent home-cages. Elife. 2021;10:e66173.

32. Nguyen KP, O’Neal TJ, Bolonduro OA, White E, Kravitz AV. Feeding Experimentation Device (FED): A flexible open-source device for measuring feeding behavior. J Neurosci Methods. 2016;267:108–114.

33. Bagley EE, Ingram SL. Endogenous opioid peptides in the descending pain modulatory circuit. Neuropharmacology. 2020;173:108131.

34. Du Y, Zhang A, Li Z, Zhao Y, Liu S, Wei C, et al. Role of astrocytic mu-opioid receptors of the ventrolateral periaqueductal gray in modulating anxiety-like responses. Behavioral and Brain Functions. 2025;21:24.

35. Godoi MM, Junior HZ, da Cunha JM, Zanoveli JM. Mu-opioid and CB1 cannabinoid receptors of the dorsal periaqueductal gray interplay in the regulation of fear response, but not antinociception. Pharmacology Biochemistry and Behavior. 2020;194:172938.

36. Huang KW, Ochandarena NE, Philson AC, Hyun M, Birnbaum JE, Cicconet M, et al. Molecular and anatomical organization of the dorsal raphe nucleus. ELife. 2019;8:e46464.

37. Okaty BW, Sturrock N, Escobedo Lozoya Y, Chang Y, Senft RA, Lyon KA, et al. A single-cell transcriptomic and anatomic atlas of mouse dorsal raphe Pet1 neurons. ELife. 2020;9:e55523.

38. Commons KG. Dorsal raphe organization. J Chem Neuroanat. 2020;110:101868.

39. Welsch L, Colantonio E, Frison M, Johnson DA, McClain SP, Mathis V, et al. Mu Opioid Receptor-Expressing Neurons in the Dorsal Raphe Nucleus Are Involved in Reward Processing and Affective Behaviors. Biol Psychiatry. 2023;94:842–851.

40. Eiden LE, Hernández VS, Jiang SZ, Zhang L. Neuropeptides and small-molecule amine transmitters: cooperative signaling in the nervous system. Cell Mol Life Sci. 2022;79:492.

41. Nusbaum MP, Blitz DM, Marder E. Functional consequences of neuropeptide and small-molecule cotransmission. Nat Rev Neurosci. 2017;18:389–403.

42. Tao R, Auerbach SB. μ-Opioids disinhibit and κ-opioids inhibit serotonin efflux in the dorsal raphe nucleus. Brain Research. 2005;1049:70–79.

43. Jolas T, Aghajanian GK. Opioids suppress spontaneous and NMDA-induced inhibitory postsynaptic currents in the dorsal raphe nucleus of the rat in vitro. Brain Res. 1997;755:229–245.

44. Xie L, Wu H, Chen Q, Xu F, Li H, Xu Q, et al. Divergent modulation of pain and anxiety by GABAergic neurons in the ventrolateral periaqueductal gray and dorsal raphe. Neuropsychopharmacol. 2023;48:1509– 1519.

45. Grahn RE, Maswood S, McQueen MB, Watkins LR, Maier SF. Opioid-dependent effects of inescapable shock on escape behavior and conditioned fear responding are mediated by the dorsal raphe nucleus. Behavioural Brain Research. 1999;99:153–167.

46. Xu Z, Feng Z, Zhao M, Sun Q, Deng L, Jia X, et al. Whole-brain connectivity atlas of glutamatergic and GABAergic neurons in the mouse dorsal and median raphe nuclei. ELife. 2021;10:e65502.

