## Supplementary Material for "Dorsal Raphe Nucleus Enkephalin Peptide Modulates Behavioral Preference"

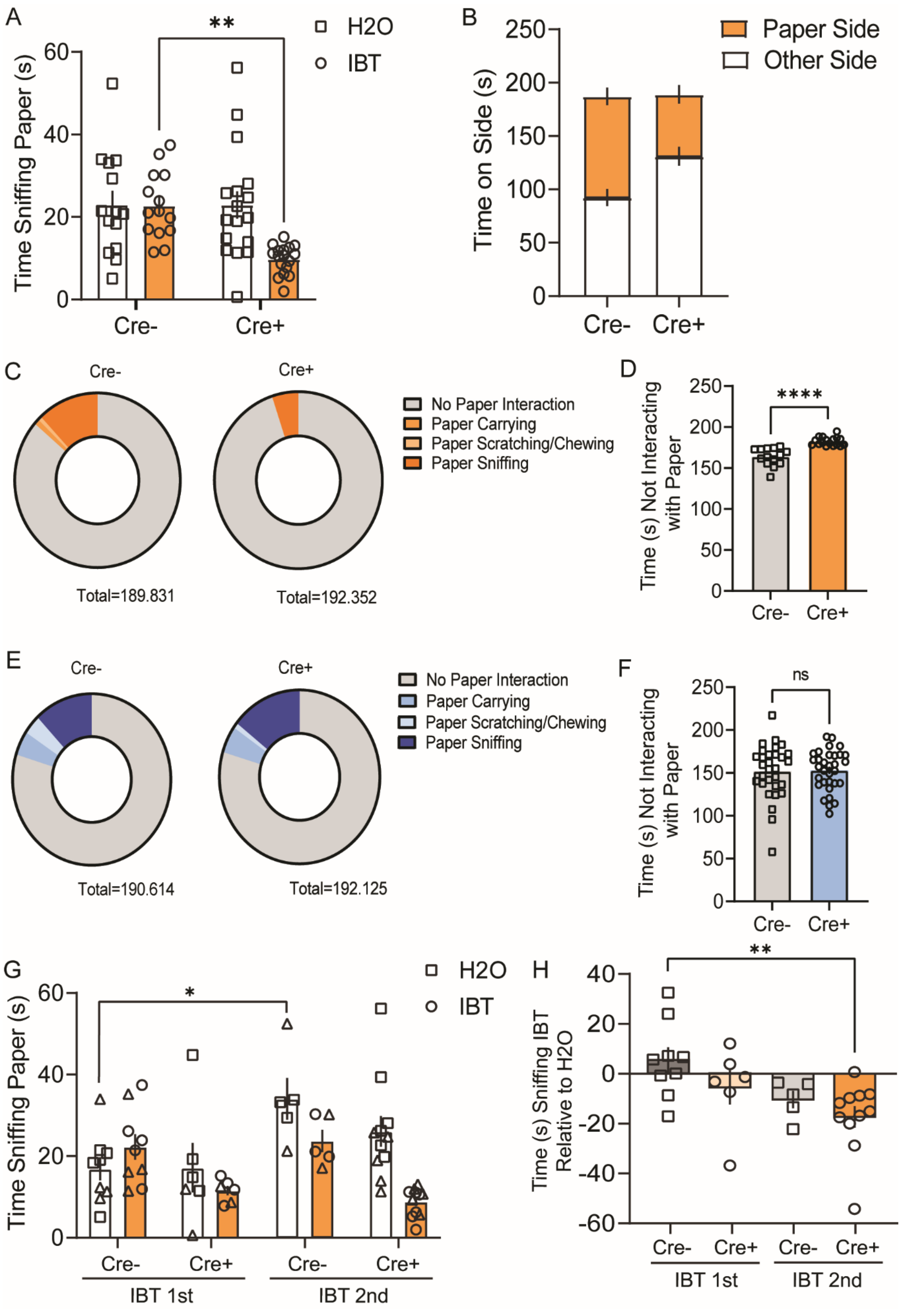


**Supplementary Figure 1: DRN$^{Penk}$ knockdown enhances aversion to IBT.** A) Raw time spent sniffing filter paper infused with either water (squares) or 2-isobutyl-thiazaole (IBT; circles) for Cre- and Cre+ mice Cre- n=14; Cre+ n=17. Cre+ mice spent less time sniffing IBT-infused filter paper than Cre- controls, Two-Way ANOVA with

Sidak's multiple comparison, **p=0.0017 B) Bar graph representing the average time during the 3-minute IBT test when the mouse was on either the side of the chamber with the paper (Paper Side; orange) or the opposite side of the chamber (Other Side; white). Cre+ mice spent more time on the Other Side than Cre- controls (Cre+ =131.0s vs Cre- =92.39s, Two-Way ANOVA with Sidak's multiple comparison, **p=0.0049) and less time on the Paper Side than Cre- controls (Cre+ =58.03s vs Cre- =94.71s, Two-Way ANOVA with Sidak's multiple comparison, **p=0.0078). C) Parts of a whole graph representing how mice interacted with the filter paper during the 3 minute IBT test with colors representing either No Paper Interaction (grey) or sniffing (dark orange), carrying (orange), or scratching/chewing (light orange) the filter paper. Cre- mice (left) spent most of the time not interacting with the paper (163.8s ± 2.874), and the time interacting was mostly spent sniffing (22.75s ± 2.178) with a small minority of time spent carrying the paper (1.876s ± 1.046) or scratching or chewing the paper (1.405s ± 1.405s). Cre+ mice (right) in contrast spent no time carrying the IBT filter paper, only a small minority scratching or chewing the paper (0.07059s ± 0.04833) and less overall time sniffing the filter paper (9.781s ± 0.8375) and more time not interacting with the paper (182.5s ± 1.257). D) Raw time scored as "Not Interacting with Paper" during the 3 min IBT test. Cre+ mice spent more time avoiding the paper than Cre- controls, Two-tailed unpaired t-test, ****p<0.0001. E) Parts of a whole graph representing how mice interacted with the filter paper during the 3 minute H2O test with colors representing either No Paper Interaction (grey) or sniffing (dark blue), carrying (blue), or scratching/chewing (light blue) the filter paper. Cre- mice (left) spent most of the time not interacting with the paper (152.0 ± 5.711), and the time interacting was spent sniffing (22.91s ± 2.147), carrying the paper (8.695s ± 3.070), and scratching or chewing the paper (7.009s ± 4.688). Cre+ mice (right) similarly spent most of the time not interacting with the paper (153.0 ± 4.140), and the time interacting was spent sniffing (27.56 ± 2.332) or carrying the paper (9.622s ± 2.243), and a small minority of time scratching or chewing the paper (1.943s ± 1.278). F) Raw time scored as "Not Interacting with Paper" during the 3 min H2O test. Cre+ mice spent the same amount of time avoiding the water paper as Cre- controls, Two-tailed unpaired t-test, p=0.8932. G) Data from A segregated by which order IBT and H2O tests were performed, with mice that were tested with IBT 1$^{st}$ on the left and mice tested with IBT 2$^{nd}$ on the right. There was an order effect, where the Cre- mice given IBT first spent less time sniffing water than those that were tested with water first, Two-Way ANOVA with Sidak's compariaon, *p= 0.0236. This effect was not as obvious in the Cre- mice p= 0.5635. H) Data from G represented as time spent sniffing IBT relative to H2O. The order effect observed with the raw times does not translate to the relative preference where each individual mouse is compared to its own "control" test of water, since there is not a significant difference between Cre- mice tested with IBT first or second, One-Way ANOVA with Tukey's multiple comparison, p= 0.2281. There are also no significant difference between the Cre+ testing order p=0.3954. But it does suggest that the order of testing drives the effect seen in Fig 1G, since the only significant difference is between Cre- mice tested with IBT 1$^{st}$ and Cre+ mice tested with IBT 2$^{nd}$ **p=0.0082. Data are all mean ± SEM.

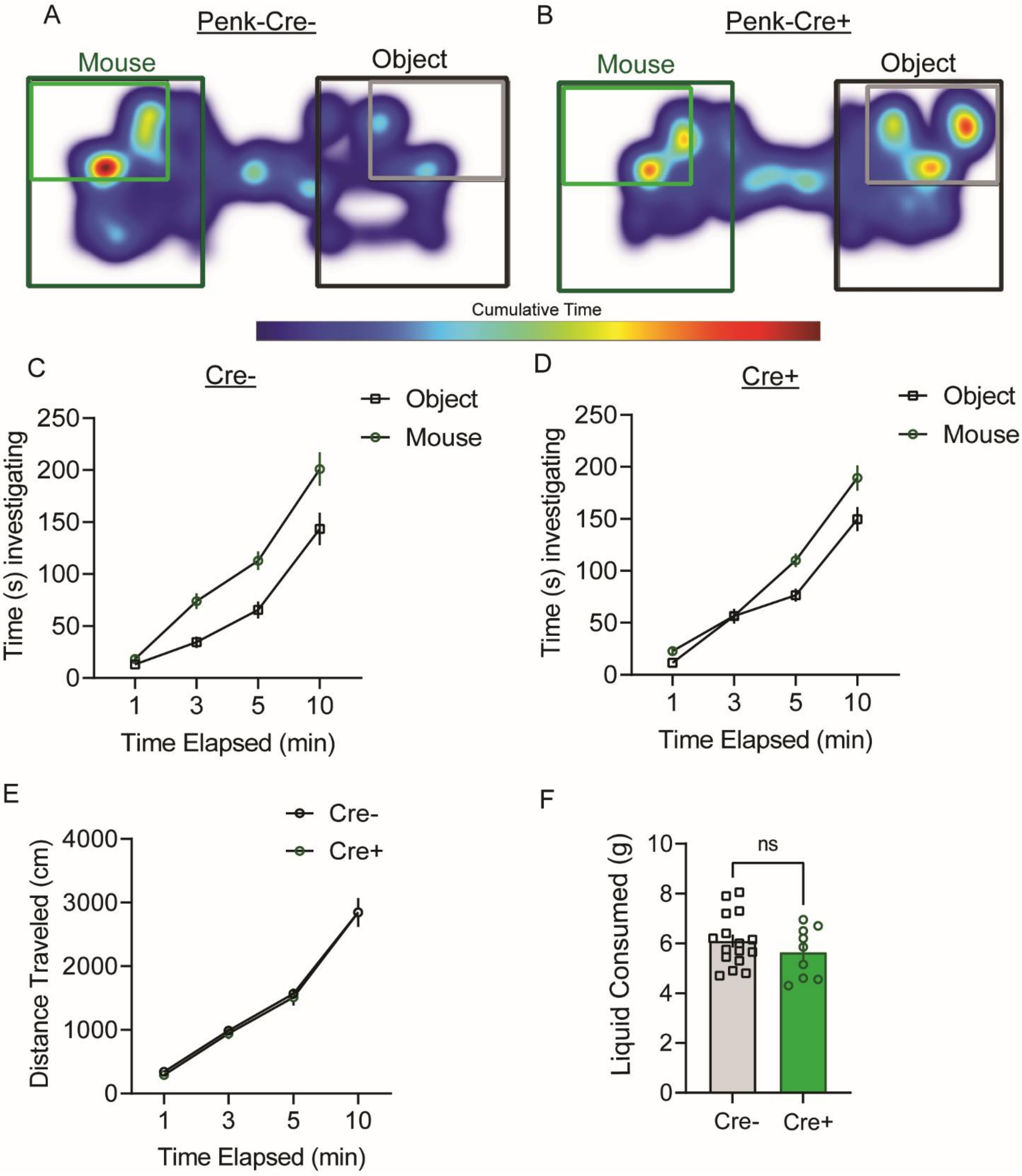


**Supplementary Figure 2: DRN$^{Penk}$ knockdown diminishes social preference but not distance moved during test or liquid consumed during sucrose preference test.** A-B) Representative heatmaps of location of Penk-Cre- (A) or Penk-Cre+ (B) mice during the first 5 minutes of a social interaction test. Green boxes represent where the stranger mouse was and black and grey boxes represent where the novel object was. Warmer colors represent more time spent in that area. C-D) Cumulative time spent in either the object (squares) or mouse (circles) innermost zones for Cre- (C) and Cre+ (D) mice. E) Cumulative distance traveled throughout the social interaction test did not differ between Cre- and Cre+ mice. F) Average liguid consumed (g) over test days in the sucrose preference test did not differ between Cre- and Cre+ mice Welch's t-test, p= 0.3114. Data are all mean ± SEM.

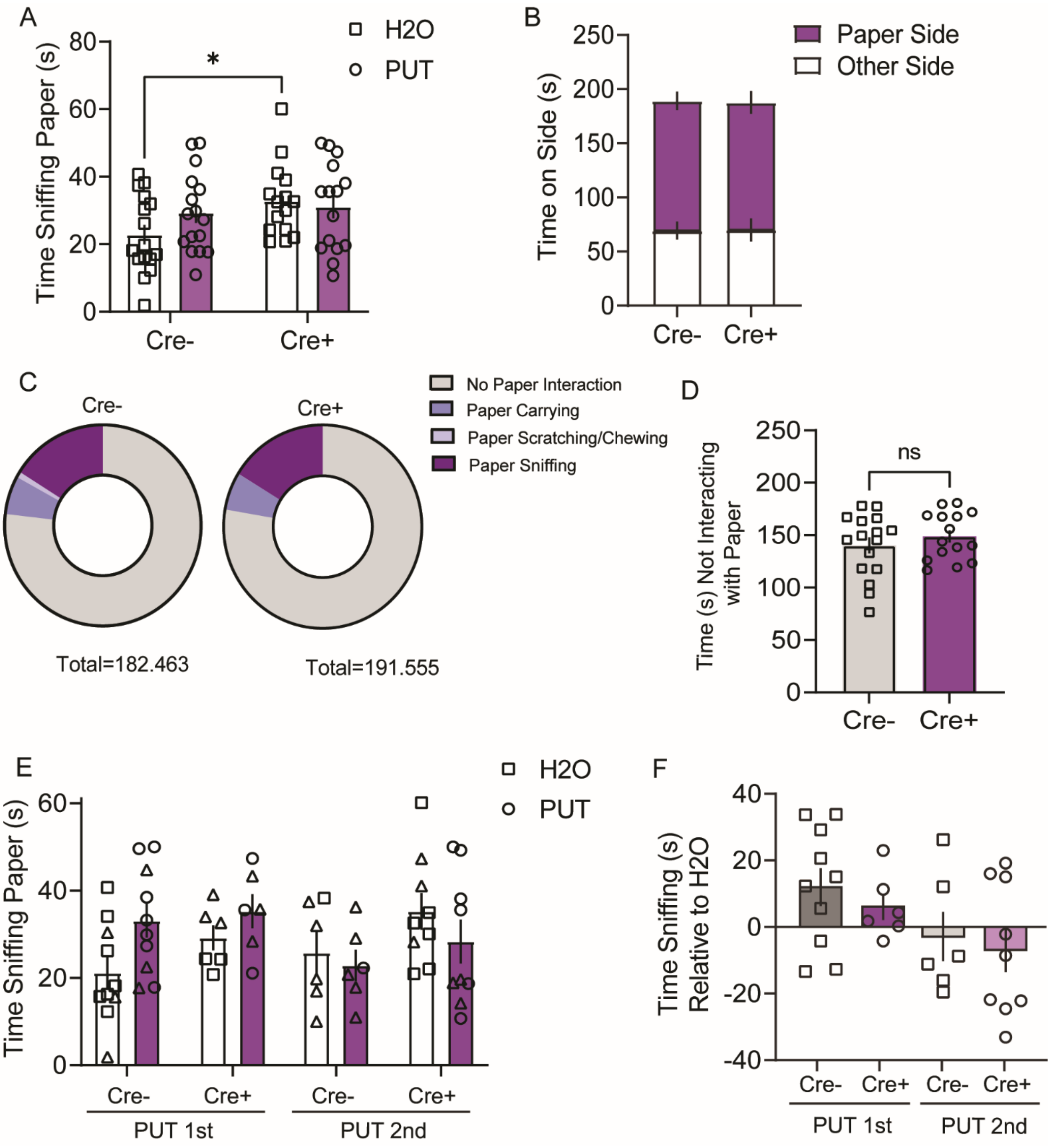


**Supplementary Figure 3: DRN$^{Penk}$ knockdown impairs preference for PUT.** A) Raw time spent sniffing filter paper infused with either water (squares) or putrescene (PUT; circles) for Cre- and Cre+ mice; Cre- n=12; Cre+ n=15. Cre+ mice spent more time sniffing H2O-infused filter paper than Cre- controls, Two-Way ANOVA with Sidak's multiple comparison, *p=0.0459 B) Bar graph representing the average time during the 3-minute PUT test when the mouse was on either the side of the chamber with the paper (Paper Side; purple) or the opposite side of the chamber (Other Side; white). Both Cre+ and Cre- mice spent more time on the Paper Side than the Other Side, Two Way ANOVA with Sidak's multiple comparison, Cre+ (Other Side=70.0 vs Paper Side= 117.7) *p=0.0232, Cre- (Other Side=69.32 vs Paper Side=119.8) *p=0.0395. C) Parts of a whole graph representing how mice interacted with the filter paper during the 3 minute PUT test with colors representing either No Paper Interaction (grey) or sniffing (dark purple), carrying (purple), or scratching/chewing (light purple) the filter paper. Cre- mice (left) spent most of the time not interacting with the paper (140.2s± 7.60), and the time interacting was spent sniffing (29.27s ± 2.981) and carrying the paper (11.25s ± 4.712), with a small amount of time spent scratching or chewing the paper (1.743s ± 1.129). Cre+ mice (right) similarly spent most of the time not interacting with the paper (148.9s ± 5.850), and the time interacting was spent sniffing (31.10s ± 3.417) or carrying the paper (11.49s ± 4.279), and a small fraction of time scratching or chewing the paper (0.06533s ± 0.6533). D) Raw time scored as "Not Interacting with Paper" during the 3 min PUT test. Cre+ mice spent the same amount of time not interacting with the PUT paper as Cre- controls, Two-tailed unpaired t-test, p=0.3725. E) Data from A segregated by which order PUT and H2O tests were performed, with mice that were tested with PUT 1$^{st}$ on the left and mice tested with PUT 2$^{nd}$ on the right. There were no significant differences between sniffing times of either H2O or PUT between the genotypes or the order of presentation, Two-Way ANOVA with Sidak's multiple comparison, H2O vs IBT; IBT 1$^{st}$ Cre- p=0.1433, Cre+ p=0.9905; IBT 2$^{nd}$ Cre- p=0.8704, Cre+ p=0.6725. F) Data from E represented as time spent sniffing PUT relative to H2O. Although it appears there is an order effect, where for either genotype the presentation of PUT 1$^{st}$ leads to a preference and the presentation of it second leads to a slight aversion, the data were not significantly different, One Way ANOVA with Tukey's multiple comparison, Cre- PUT 1$^{st}$ vs Cre+ PUT 2$^{nd}$, p=0.1079. Data are all mean ± SEM.

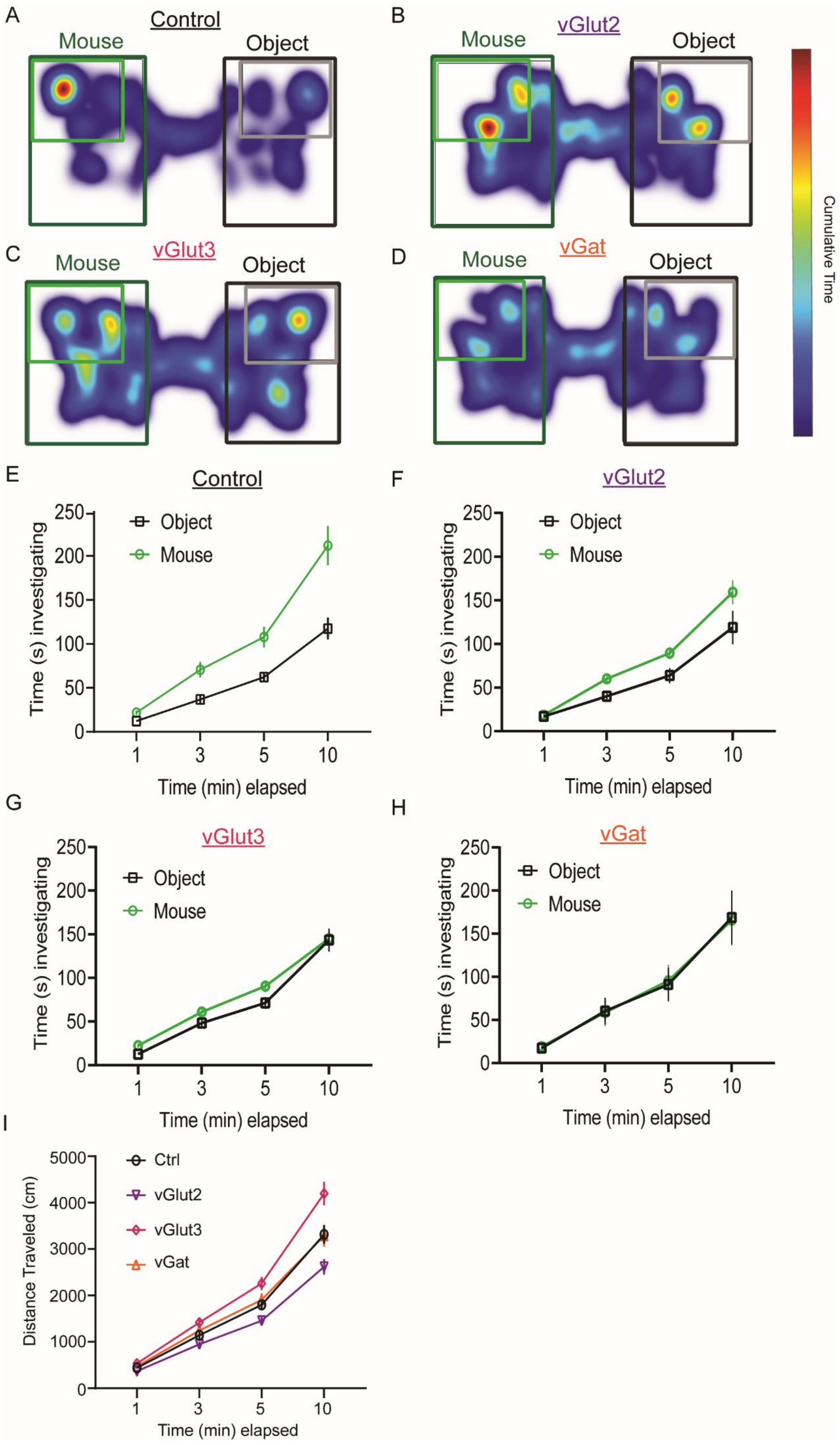
A
Control
Mouse
Object
B
vGlut2
Mouse
Object
Cumulative Time
C
vGlut3
Mouse
Object
D
vGat
Mouse
Object
E
Control
Object
Mouse
Time (s) investigating
Time (min) elapsed
F
vGlut2
Object
Mouse
Time (s) investigating
Time (min) elapsed
G
vGlut3
Object
Mouse
Time (s) investigating
Time (min) elapsed
H
vGat
Object
Mouse
Time (s) investigating
Time (min) elapsed
I
Ctrl
vGlut2
vGlut3
vGat
Distance Traveled (cm)
Time (min) elapsed

**Supplementary Figure 4: Social Interaction effects of DRN$^{Penk}$ knockdown does not depend on glutamatergic or GABAergic cell populations alone.** A-D) Representative heatmaps of location of Control (A), vGlut2 (B), vGlut3 (C), or vGat (D) mice during the 10 minutes of a social interaction test. Green boxes represent where the stranger mouse was and black and grey boxes represent where the novel object was. Warmer colors represent more time spent in that area. E-H) Cumulative time spent in either the object (squares) or mouse (circles) innermost zones for Control (E), vGlut2 (F), vGlut3 (G), or vGat (H) mice. I) Cumulative distance traveled throughout the social interaction test did not differ between vG-Cre and Control mice (Two Way ANOVA with Tukey's multiple comparison, $p>0.05$). Data are all mean ± SEM.

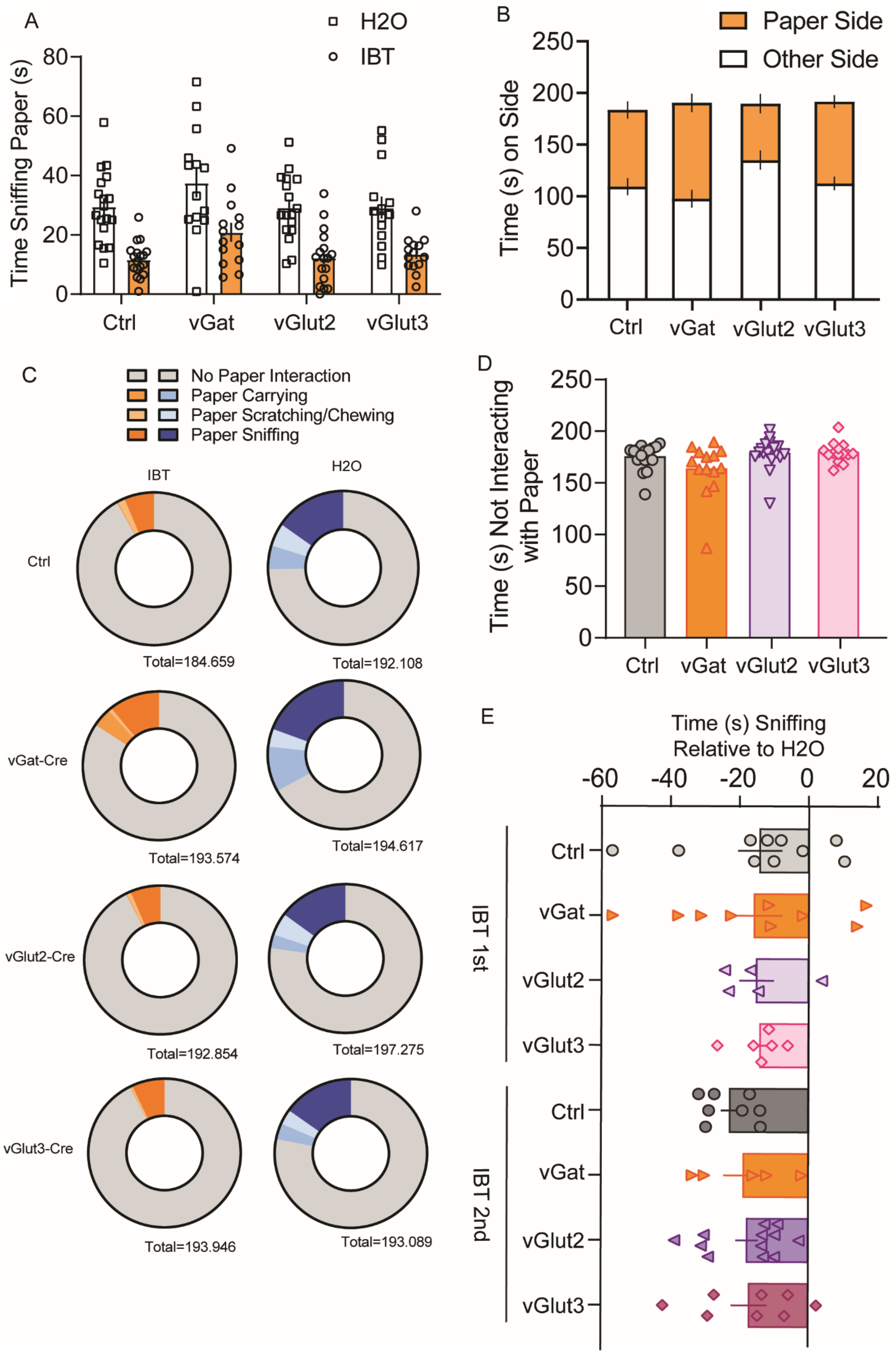

A
H2O
IBT
Time Sniffing Paper (s)
Ctrl
vGat
vGlut2
vGlut3
B
Paper Side
Other Side
Time (s) on Side
Ctrl
vGat
vGlut2
vGlut3
C
No Paper Interaction
Paper Carrying
Paper Scratching/Chewing
Paper Sniffing
IBT
H2O
Ctrl
Total=184.659
Total=192.108
vGat-Cre
Total=193.574
Total=194.617
vGlut2-Cre
Total=192.854
Total=197.275
vGlut3-Cre
Total=193.946
Total=193.089
D
Time (s) Not Interacting with Paper
Ctrl
vGat
vGlut2
vGlut3
E
Time (s) Sniffing Relative to H2O
IBT 1st
IBT 2nd
Ctrl
vGat
vGlut2
vGlut3

**Supplementary Figure 5: IBT Aversion effects of DRN$^{Penk}$ knockdown are not dependent on glutamatergic or GABAergic subpopulations.** A) Raw time spent sniffing filter paper infused with either water (squares) or 2-isobutyl-thiazaole (IBT; circles) for Control and vG-Cre mice Contol n= 18, vGlut2 n= 17, vGlut3 n= 14, vGat n= 14. B) Bar graph representing the average time during the 3-minute IBT test when the mouse was on either the side of the chamber with the paper (Paper Side; orange) or the opposite side of the chamber (Other Side; white). None of the vG-Cre subtypes spent more or less time on the Other Side than controls (Ctrl =108.1s vs vGat =97.63, vGlut2 =135.0s, vGlut3 =112.4s, Two-Way ANOVA with Sidak's multiple comparison, $p>0.05$) and the same was true for the Paper Side. C) Parts of a whole graph representing how mice interacted with the filter paper during the 3 minute IBT (left; orange hues) or H2O (right; blue hues) tests with colors representing either No Paper Interaction (grey) or sniffing (dark orange/blue), carrying (orange/blue), or scratching/chewing (light orange/blue) the filter paper. D) Raw time scored as "Not Interacting with Paper" during the 3 min IBT test. Control and vG-Cre mice spent the same amount of time avoiding the filter paper, One Way ANOVA with Tukey's multiple comparison, $p>0.05$ E). Data from A segregated by which order IBT and H2O tests were performed and normalized by the equation (Time sniffing IBT- Time Sniffing H2O), with mice that were tested with IBT 1$^{st}$ on the top and mice tested with IBT 2$^{nd}$ on the bottom. There were no significant differences between genotypes within testing order (One way ANOVA with Tukey's multiple comparison, IBT 1$^{st}$/2$^{nd}$ Ctrl vs vG-Cre's, $p>0.05$) and no significant differences within genotypes between the order of testing (One way ANOVA with Tukey's multiple comparison, Ctrl/vG-Cre IBT 1$^{st}$ vs Ctrl/vG-Cre IBT 2$^{nd}$, $p>0.05$). Data are all mean ± SEM.
